## Supplementary Information for "Courtship behaviour reveals temporal regularity is a critical social cue in mouse communication"

### Supporting information

#### Audio files S1. Audible samples of sounds used in the playback experiments

A. Intact song (top panel in Fig. 3A)

B. Randomized song (bottom panel in Fig. 3A)

C. Reversed song (bottom panel in Fig. 3C)

D. Phase-scrambled song (bottom panel in Fig. 3E)

E. Pure tone syllables (bottom panel in Fig. 3G)

F. Irregular song (bottom panel in Fig. 4E)

G. Super-regular song (bottom panel in Fig. 4H)

#### Video file S2. Example video tracking of mouse approach behaviour

A. Third sound presentation trial from example behavioural session (green trace in Fig. 2C), with intact song playback from the left speaker.

B. Fourth sound presentation trial from example behavioural session (green trace in Fig. 2C), with intact song playback from the left speaker.

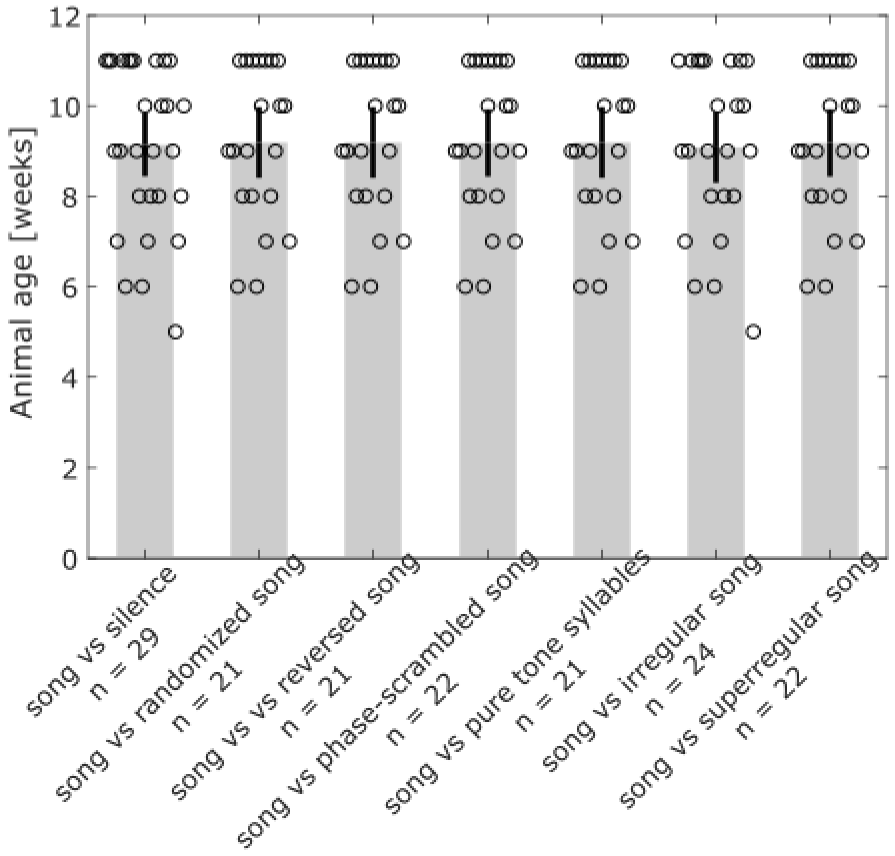


**Figure S3. No difference in listener age across playback experiments**

The mice participating in each type of playback experiment were of similar age (one-way ANOVA, $F(6, 153) = 0.013$, $p = 1.0$). Shown is mean and 95% confidence interval.

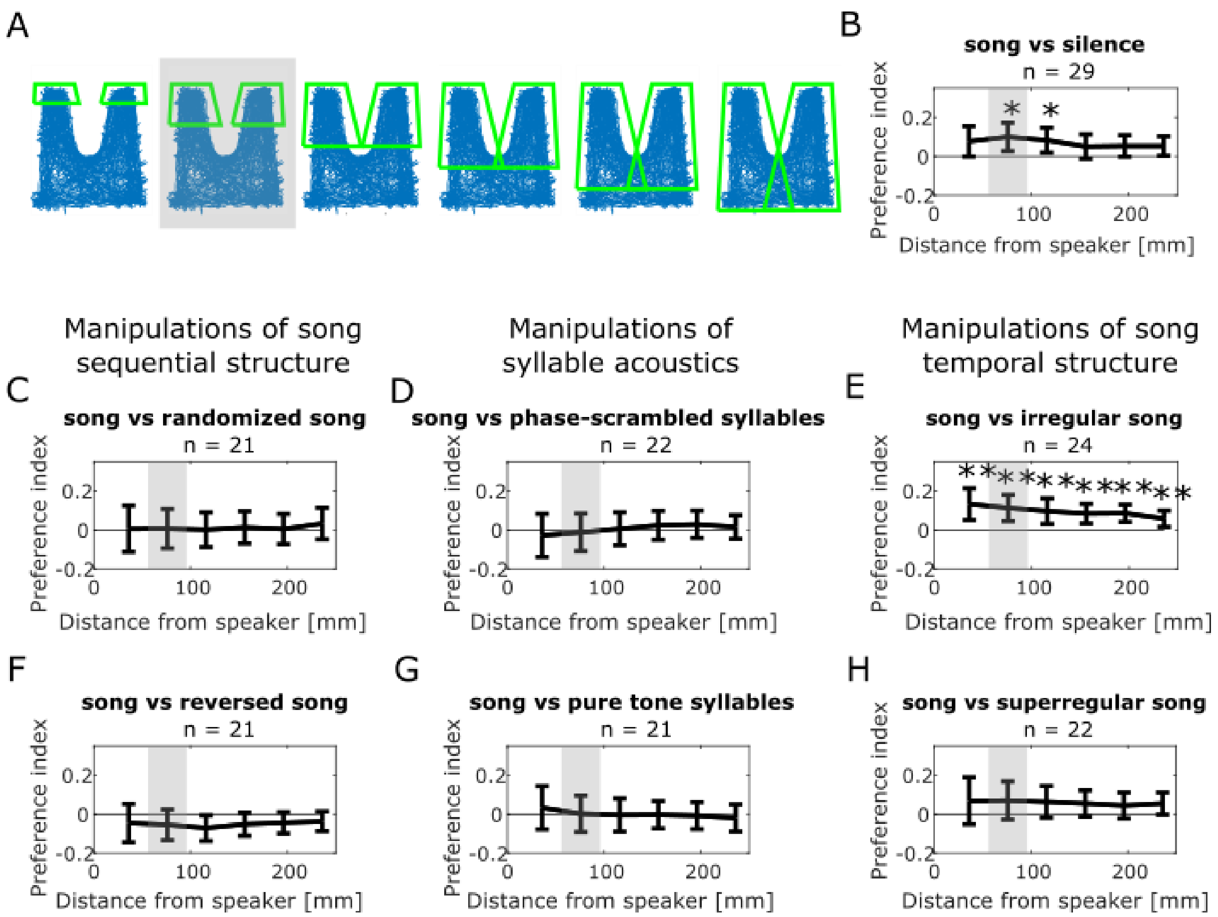


**Figure S4. Approach behaviour is consistent across varying speaker zone lengths**

A. Tracking of the mouse's position over the course of a behavioural session, in the two-compartment behavioural box with a soundproof partition (middle) positioned between two ultrasonic loudspeakers (top). Green outlines show the two "speaker zones" used as regions of interest for quantifying the animal's position. The grey-shaded image illustrates the speaker zone used throughout the study. Additional images illustrate speaker zones with varying lengths, as the distal edge is moved closer and/or further away from the speakers.

B. Population preference index used to quantify approach behaviour in response to playback of intact songs (positive values) relative to silence (negative value), for varying speaker zone lengths in panel A. The grey shading indicates the speaker zone used throughout the study. Shown is mean and 95% confidence interval. One-sample two-tailed *t*-test, **: $p < 0.01$, ns: non significant, $p > 0.05$

C. Population preference index used to quantify approach behaviour in response to intact songs (positive values) relative to sequences of randomly ordered syllables (negative value), displayed as in panel B.

D. Population preference index used to quantify approach behaviour in response to intact songs (positive values) relative to sequences of phase-scrambled syllables (negative value).

E. Population preference index used to quantify approach behaviour in response to intact (positive values) relative to temporally irregular (negative value) songs.

F. Population preference index used to quantify approach behaviour in response to intact (positive values) relative to reversed (negative value) songs.

G. Population preference index used to quantify approach behaviour in response to intact songs (positive values) relative to sequences of pure-tone approximation of syllables (negative value).

H. Population preference index used to quantify approach behaviour in response to intact (positive values) relative to temporally superregular (negative value) songs.

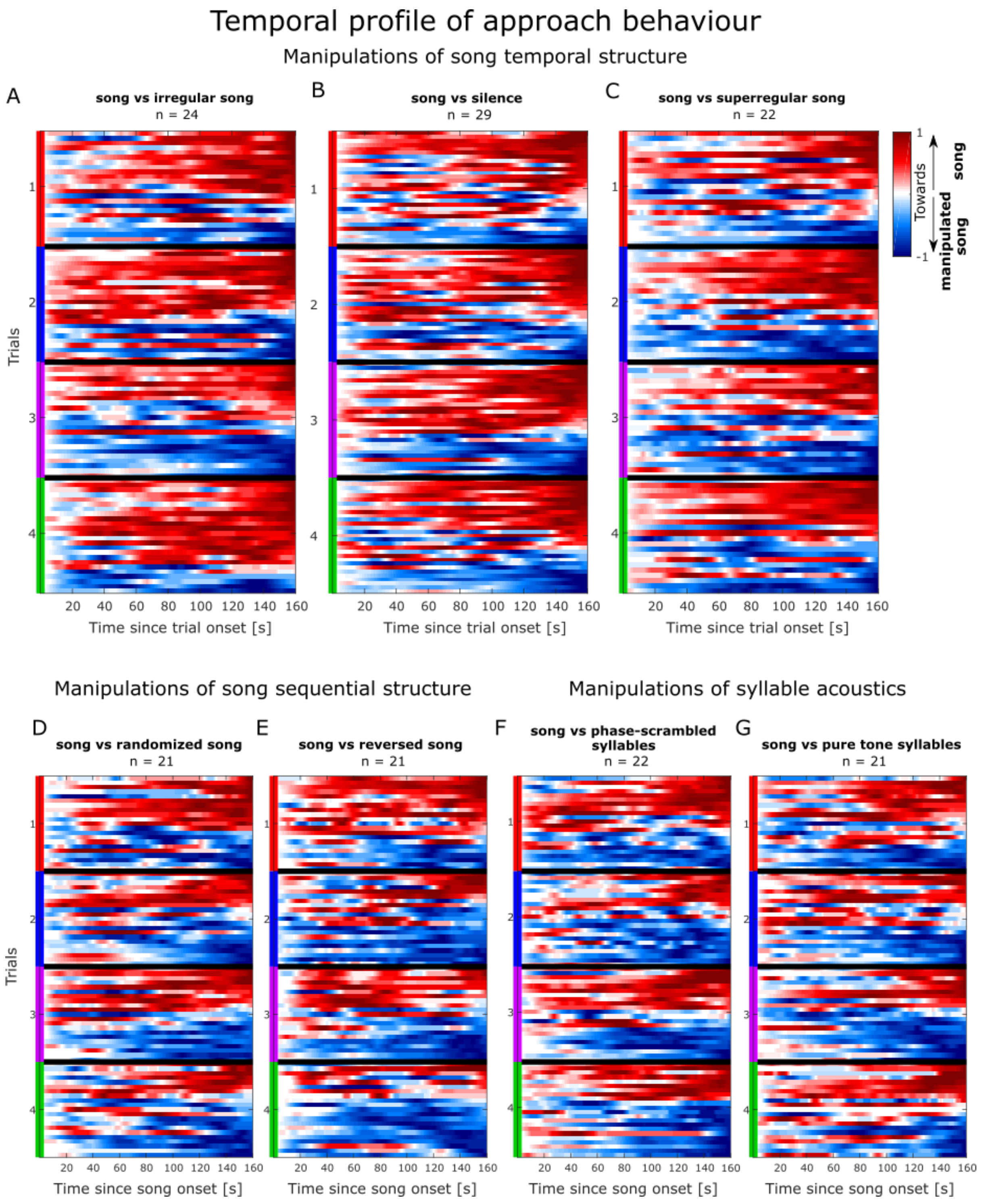


**Figure S5: Trial-based temporal profiles of approach behaviour.**

A Timecourse of approach behaviour during one presentation of intact (positively weighted) vs temporally irregular (negatively weighted) mouse song (see Fig. 2E), grouped by sound presentation trial (coloured bars), for each behavioural session (n = 24, y-axis). Each trial's trace was normalized to the absolute value of its maximal amplitude. Within a trial, sessions are ordered by the amplitude of their last element.

B. Timecourse of approach behaviour during one presentation of intact mouse songs (positively weighted) vs silence (negatively weighted).

C. Timecourse of approach behaviour during one presentation of intact (positively weighted) vs temporally superregular (negatively weighted) mouse song.

D. Timecourse of approach behaviour during one presentation of intact mouse song (positively weighted) vs sequences of randomly ordered syllables (negatively weighted).

E. Timecourse of approach behaviour during one presentation of intact (positively weighted) vs reversed (negatively weighted) mouse song

F. Timecourse of approach behaviour during one presentation of intact mouse songs (positively weighted) vs sequences of phase-scrambled syllables (negatively weighted).

G. Timecourse of approach behaviour during one presentation of intact mouse songs (positively weighte d) vs sequences of pure-tone approximation of syllables (negatively weighted).
